# Ovine MyD88 have active role in host resistance against *Haemonchus contortus* infection

**DOI:** 10.1101/2023.10.08.561398

**Authors:** Samiddha Banerjee, Aruna Pal, Abantika Pal, Jayanta Chatterjee

## Abstract

Host resistance against parasitic infestation is quite important to study in current days, since till date no effective vaccines against parasites have been developed. Moreover, indiscrimate use of regular deworming as a practice may lead to the development of resistance to antihelmiths (as in case of antibiotic resistance against bacteria) or development of antihelminth resistant strains among dreadly parasites. In this present study, we attempt to explore immune response gene against *Haemonchus contortus* as dreadly parasite against sheep model. In our lab, we have already detected some of such promising genes, namely RIGI (earlier known antiviral in action), CD14, IL6 and IL10 (earlier known antibacterial in action). In this current publication, we have explored the role of MyD88 in mediating resistance against parasitic infestation. We have characterized ovine MyD88 molecule, identified important domain, analyzed molecular docking, identified the binding site with *H. contortus* through *in silico* studies, network and pathway analysis, followed by differential mRNA expression profiling, ultimate confirmation through immunohistochemistry. Identification of immune response genes against parasites may lead to immunomodulation, genomic selection, gene therapy or even development of disease resistant stock of sheep through gene editing approach.

## Introduction

Host resistance against parasitic infestation is a novel area to study, since till date very scanty studies have been undertaken unlike bacterial or viral infestation. Antiparasitic drugs, including antihelminths are used for treatment of parasities, endoparasites.Gastrointestinal parasites like *Haemonchus contortus* is one of the most dreadly parasites, causing heavy economic loss in sheep industry. Control of Haemonchus contortus infestation is becoming exceedingly difficult. So far we routinely administer anihelminthics as routine farm management practice and also advocate deworming procedure to sheep/goat maintained by farmers for both prophylaxis and also for treatment. This may cause resistance to particular parasite, like development of antibiotic resistance to bacteria. Moreover, another limitation is the lack of effective approved vaccine against Haemonchus contortus. An attempt has been made with **Barbervax**^®^ with approval in limited areas (https://wormboss.com.au). Moreover they claim immunity to lasts for only six weeks, which is too costly and of not much effective. As a result, the best option is to control through immunomodulation. In our earlier studies, we have studied immune response genes RIGI, CD14, IL6 and IL10 to have important role in providing resistance against *Haemonchus contortus (*Banerjee et al., 2021, Rawat et al., 2021, Rawat et al., 2023).

The mechanism of action of MyD88 against bacteria has been described by some researchers. MyD88 (Myeloid Differentiation Primary Response gene 88) transfers signals from certain proteins called Toll-like receptors and interleukin-1 (IL-1) receptors, which are important for an early immune response to foreign invaders such as bacteria. In response to signals from these receptors, the MyD88 adapter protein stimulates signaling molecules that turn on a group of interacting proteins which are nuclear factor-kappa-B. It is an important adaptor protein which is used by every Toll like receptor except TLR3. TIRAP (Toll/Interleukin-1 receptor domain containing adaptor protein) is required for the recruitment of MyD88 to TLR-2 and TLR-4 and then it plays its part and signalling occurs through IRAK (Interleukin-1 receptor associated kinase) (Arancibia SA et.al. 2007.). The MYD88 gene instructs to make a protein engaged in signalling in the immune cells. The MyD88 protein acts as an adapter, connecting proteins that receive signals from outside the cell to the proteins that relay signals inside the cell.

In our laboratory, we have already explored a set of genes with the role in resistance against Haemonchus contortus, which were earlier known as antiviral (RIGI) or antibacterial (CD14, IL6, IL10) molecule.

Considering the above studies, it had been observed that MyD88 known well as providing antibacterial immunity. Very scanty report for MyD88 are available for immune function against parasites. Hence the present study was undertaken for molecular characterization of MyD88 of sheep, differential mRNA expression profile for MyD88 gene in healthy and infected sheep with respect to *Haemonchus contortus* infection, and predicting the binding site with H.contortus for these molecules.

## Materials and Method

### Animals and RNA isolation

The present study was conducted on Garole sheep maintained in Livestock Farm Complex, West Bengal University of Animal and fishery Sciences. All the animals were maintained under uniform managemental conditions and are of similar age group (1-2 years). Total RNA was isolated from abomassum of sheep, which is actually the site of predilection for Haemonchus contortus following Trizol method (Pal et al, 2011, Pal et al., 2014, Pal et al., 2021).

### Materials

Taq DNA polymerase, 10X buffer, dNTP were purchased from Invitrogen, SYBR Green qPCR Master Mix (2X) was obtained from Thermo Fisher Scientific Inc. (PA, USA). Unlabelled goat anti-bovine polyclonal antibodies against MyD88 were obtained from Santa Cruz Biotechnology Inc, (CA, USA) and horseradish peroxidase-labeled rabbit anti-goat antibody was obtained from Sigma-Aldrich (St. Louis, MO, USA). Poly (I:C)/LyoVec HMW (endotoxin level<0.001 EU/µg) was obtained from Invivogen (San Diego, CA). Trypsin-EDTA, foetal bovine serum (FBS) and Epidermal Growth Factor (EGF) were obtained from Sigma-Aldrich (St. Louis, MO, USA). L-Glutamine (Glutamax 100x) was purchased from Invitrogen corp., (Carlsbad, CA, USA). Penicillin-G and streptomycin were obtained from Amresco (Solon, OH, USA). Filters (Millex GV. 0.22 µm) were purchased from Millipore Pvt. Ltd., (Billerica, MA, USA). All other reagents were of analytical grade.

The 20 μL reaction mixture contained 5 μg of total RNA, 0.5 μg of oligo dT primer (16–18Lmer), 40LU of Ribonuclease inhibitor, 1000LM of dNTP mix, 10 mM of DTT, and 5LU of MuMLV reverse transcriptase in reverse transcriptase buffer. The reaction mixture was gently mixed and incubated at 37°C for 1 hour. The reaction was stopped by heating the mixture at 70°C for 10 minutes and chilled on ice. The integrity of the cDNA checked by PCR. The primers were visualized in Table. 25 μL reaction mixture contained 80–100 ng cDNA, 3.0 μL 10X PCR assay buffer, 0.5 μL of 10 mM dNTP, 1LU Taq DNA polymerase, 60 ng of each primer, and 2LmM MgCl2. PCR-reactions were carried out in a thermocycler (PTC-200, MJ Research, USA) with cycling conditions as, initial denaturation at 94°C for 3 min, denaturation at 94°C for 30Lsec, annealing at 60°C for 35Lsec, and extension at 72°C for 3 min were carried out for 35 cycles followed by final extension at 72°C for 10Lmin.

### 2.3. cDNA Cloning and Sequencing

PCR amplicons verified by 1% agarose gel electrophoresis were purified from gel using Gel extraction kit (Qiagen GmbH, Hilden, Germany) and ligated into pGEM-T easy cloning vector (Promega, Madison, WI, USA) following manufacturers’ instructions. The 10 μL of ligated product was directly added to 200 μL competent cells, and heat shock was given at 42°C for 45Lsec. in a water bath, and cells were then immediately transferred on chilled ice for 5 min., and SOC was added. The bacterial culture was pelleted and plated on LB agar plate containing Ampicillin (100 mg/mL) added to agar plate @ 1L:L1000, IPTG (200 mg/mL) and X-Gal (20 mg/mL) for blue-white screening. Plasmid isolation from overnight-grown culture was done by small-scale alkaline lysis method (Pal et al., 2021, Sahu et al,2023a, Sahu et al,2023b, Sahu et al,2023c). Recombinant plasmids were characterized by PCR using gene-specific primers and restriction enzyme digestion based on reported nucleotide sequence for cattle. The enzyme EcoR I (MBI Fermentas, USA) isused for fragment release.Gene fragment insert in recombinant plasmid was sequenced by automated sequencer (ABI prism) using dideoxy chain termination method with T7 and SP6 primers (Chromous Biotech, Bangalore).

### 2.4. Sequence Analysis

The nucleotide sequence so obtained was analyzed for protein translation, sequence alignments, and contigs comparisons by DNASTAR Version 4.0, Inc., USA. Novel sequence was submitted to the NCBI Genbank and accession number MK986727.1 was obtained which is available in public domain now.

### 2.5. Study of Predicted MyD88 peptide Using Bioinformatics Tools

Predicted peptide sequence of MyD88 peptide of Garole sheep was derived by Edit sequence (Lasergene Software, DNASTAR) and then aligned with the peptide of other sheep breed Megalign sequence Programme of Lasergene Software (DNASTAR). Prediction of signal peptide of MyD88 gene was conducted using the software (Signal P 3.0 Sewer-prediction results, Technical University of Denmark). Leucine percentage was calculated manually from predicted peptide sequence. Di-sulphide bonds were predicted using suitable software (http://bioinformatics.bc.edu/clotelab/DiANNA/) and by homology search with other species.

Protein sequence level analysis study was carried out with specific software (http://www.expasy.org./tools/blast/) for determination of leucine rich repeats (LRR), leucine zipper, N-linked glycosylation sites, detection of Leucine-rich nuclear export signals (NES), and detection of the position of GPI anchor. Detection of Leucine-rich nuclear export signals (NES) was carried out with NetNES 1.1 Server, Technical university of Denmark. Analysis of O-linked glycosylation sites was carried out using NetOGlyc 3.1 server (http://www.expassy.org/), whereas N-linked glycosylation site were detected by NetNGlyc 1.0 software (http://www.expassy.org/). Regions for alpha helix and beta sheet were predicted using NetSurfP-Protein Surface Accessibility and Secondary Structure Predictions, Technical University of Denmark (Glick, 1977). Domain linker prediction was done according to the software developed (Ebina et al., 2009). LPS-binding (Cunningham et al., 2000) and LPS-signalling sites (Muroi et al., 2002) were predicted based on homology studies with other species MyD88 polypeptide.

### 2.6 Three dimensional structure prediction and Model quality assessment

The templates which possessed highest sequence identity with our target template were identified by using PSI-BLAST (http://blast.ncbi.nlm.nih.gov/Blast). The homology modelling was used to build 3D structure based on homologous template structures using PHYRE2 server. The 3D structures were visualized by PyMOL (http://www.pymol.org/) which is an open source molecular visualization tool. Subsequently, the mutant model was generated using PyMoL tool. The Swiss PDB Viewer was employed for controlling energy minimization. The structural evaluation along with stereo-chemical quality assessment of predicted model were carried out by using the SAVES (Structural Analysis and Verification Server), which is integrated server (http://nihserver.mbi.ucla.edu/SAVES/). The ProSA (Protein Structure Analysis) web server (https://prosa.services.came.sbg.ac.at/prosa) was used for refinement and validation of protein structure. The ProSA was used for checking model structural quality with potential errors and program shows a plot of its residue energies and Z-scores which determine overall quality of model. The solvent accessibility surface area of the IR genes was generated by using NetSurfP server (http://www.cbs.dtu.dk/services/NetSurfP/). It calculates relative surface accessibility, Z-fit score, probability for Alpha-Helix, probability for beta-strand and coil score, etc. TM align soft ware was used for alignment of 3 D structure of IR protein for different species and RMSD estimation to assess the structural differentiation. I-mutant analysis was conducted for mutations detected to assess the thermodynamic stability. Provean analysis was conducted to assess the deleterious nature of the mutant amino acid.

### 2.4. Protein-protein interaction network depiction

In order to understand network of MyD88 with related peptide, we performed analysis with submitting FASTA sequences to STRING 9.1. In STRING, the functional interaction was analyzed by using confidence score. Interactions with score < 0.3 are considered as low confidence, scores ranging from 0.3 to 0.7 are classified as medium confidence and scores > 0.7 yield high confidence. The functional partners were depicted.

KEGG analysis also depicts the functional association of MyD88 peptide with other related proteins.

### PCR Amplification

All PCR amplifications were performed in 25 µl reaction volume. Each reaction contained 2.5 µl 10X buffer, 200 µM of dNTPs, 0.5 µl of each primers (10 pmol), 0.5 units of Taq DNA polymerase and nuclease free water to bring the total volume to 25 µl. Around 100 ng of cDNA was used as template. The details of primers for different fragments of GH gene are summarized in Table 1. Thermal cycling was performed on an ABI PCR machine where a touchdown protocol with annealing 46°C for 10 cycles followed by 48 °C for 25 cycles was followed for 428 bp fragment. The PCR products were resolved on a 1.5% agarose gel (Figure S1 in <u>File S1</u>). The PCR products were cloned on to pGEMT vector, plasmids were screened using PCR, and positive plasmids were custom sequenced. cDNA was amplified for complete cds region of growth hormone gene for two different variants of Growth hormone gene.

**Table 1:**
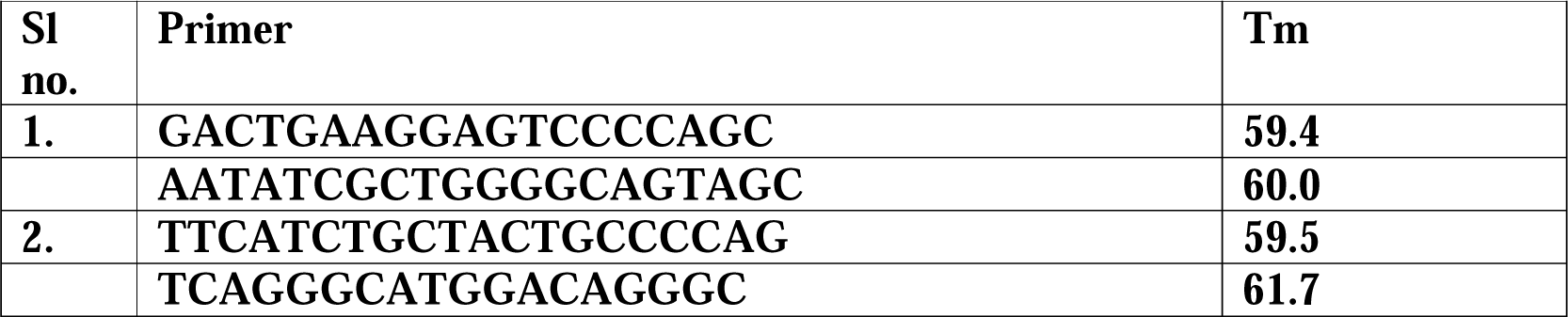
Primer sequences for amplification of MyD88 gene for sheep.

### Real Time PCR (qRT-PCR)

Real time PCR primers were designed by aligning gene sequences of several mammals including cow, pig, mouse, buffalo, and human (Table 2). QPCR study was conducted from cDNA derived from different organs collected from slaughter house during the process of slaughter. Equal amount of RNA (quantified by Qubit fluorometer, Invitrogen), wherever applicable, were used for cDNA preparation (Superscript III cDNA synthesis kit; Invitrogen). All qRT-PCR reactions were conducted on ABI 7500 fast system. Each reaction consisted of 2 µl cDNA template, 5 µl of 2X SYBR Green PCR Master Mix, 0.25 µl each of forward and reverse primers (10 pmol/µl) and nuclease free water for a final volume of 10 µl. Each sample was run in duplicate. Analysis of real-time PCR (qRT-PCR) was performed by delta-delta-Ct (ΔΔCt) method <u>[Pal et al 2021]</u>.

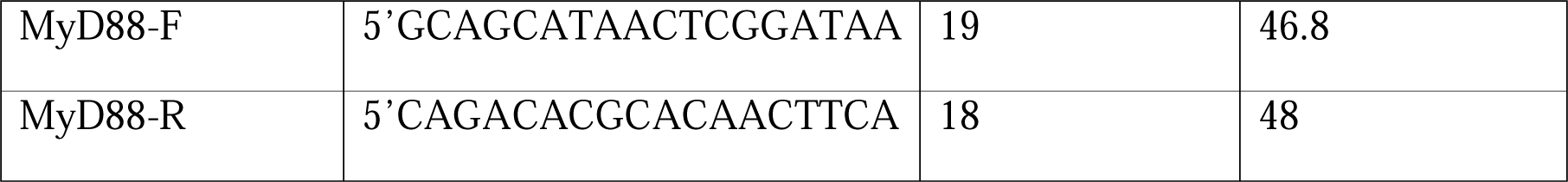

18S rRNA primer: F-TCCAGCCTTCCTTCCTGGGCAT,

R-GGACAGCACCGTGTTGGCGTAGA.

### Molecular Docking

Molecular docking is a bioinformatics tool used for *in silico* analysis for the prediction of binding mode of a ligand with a protein 3D structure. Patch dock is an algorithm for molecular docking based on shape complementarity principle (Schneidman-Duhovny et al., 2005). Patch Dock algorithm was used to predict ligand protein docking for surface antigen for *Haemonchus contortus* (alpha tubulin and beta tubulin) with MyD88. TM align was used for alignment of 3D structure for two molecules.

## Results and Discussion

### Molecular characterization and domain analysis of MyD88 in sheep

3D structural analysis for MyD88 molecule in sheep is being depicted in Fig 1, with TIR (Toll interleukin receptor) domain 1-88, 159-293. The **toll-interleukin-1 receptor (TIR) homology domain** is an intracellular cytoplasmic signaling domain found in MyD88(*Horsefield et al.,2019, Essuman et al., 2018*), It contains three highly conserved regions, and mediates protein-protein interactions between the toll-like receptors (TLRs) and signal-transduction components. When activated, TIR domains recruit cytoplasmic adaptor proteins MyD88 The intermediate domain (ID) is required for the phosphorylation and activation of IRAK at aa position 110-115. Death domain at aa position 32-109. Apoptotic signals are transduced by five death domain (Singh et al., 1998)-containing receptors--TNFR1, Fas, DR3, DR4, and DR5--by binding to their ligands. The intracellular portion of all these receptors contains a region, approximately 80 amino acids long, referred to as the “death domain” (DD).

**Fig 1:**
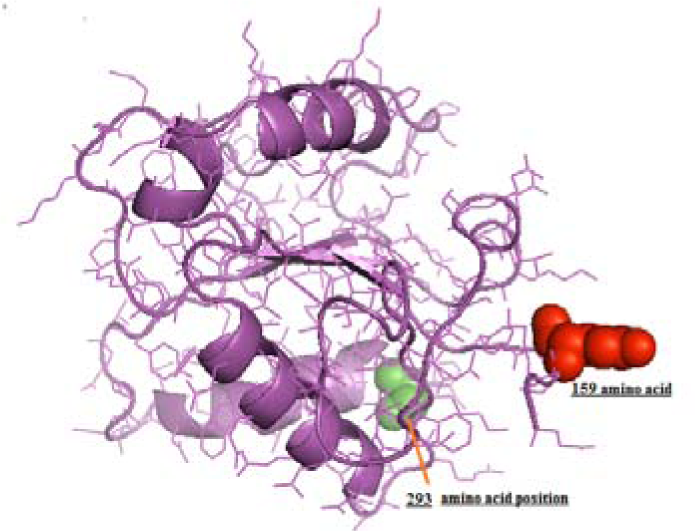
TIR domain for MyD88 from amino acid position 159 to 293aa.

### Differential mRNA expression profile of **MyD88** gene in sheep

Differential expression profiling for MyD88 indicates better expression profile for sheep infected with *Haemonchus contortus* compared to heallthy sheep. TLR4 activates the MyD88- and TRIF-dependent pathways through nuclear factor-κB (NF-κB)–associated signaling events, It triggers tumor necrosis factor alpha (TNF-α) and it is associated with the activation of the pro-inflammatory cytokines that cause the inflammatory cascade reaction (Lu *et al*., 2008).

### Molecular docking for the detection of binding site of MyD88 with respect to Haemonchus contortus

Patchdock score for two surface protein for Haemonchus contortus (alpha tubulin and beta tubulin) with MyD88 (Supplementary Fig 1 &2) was found to be less, it indicates it is not a receptor and do not bind to the parasite *Haemonchus contortus*. Hence, further binding site identification has not been processed.

### STRING Analysis/ Network analysis

STRING network analysis depicts the molecular interaction of MyD88 with other molecules of innate immune importance (Fig 3).

**Fig 2:**
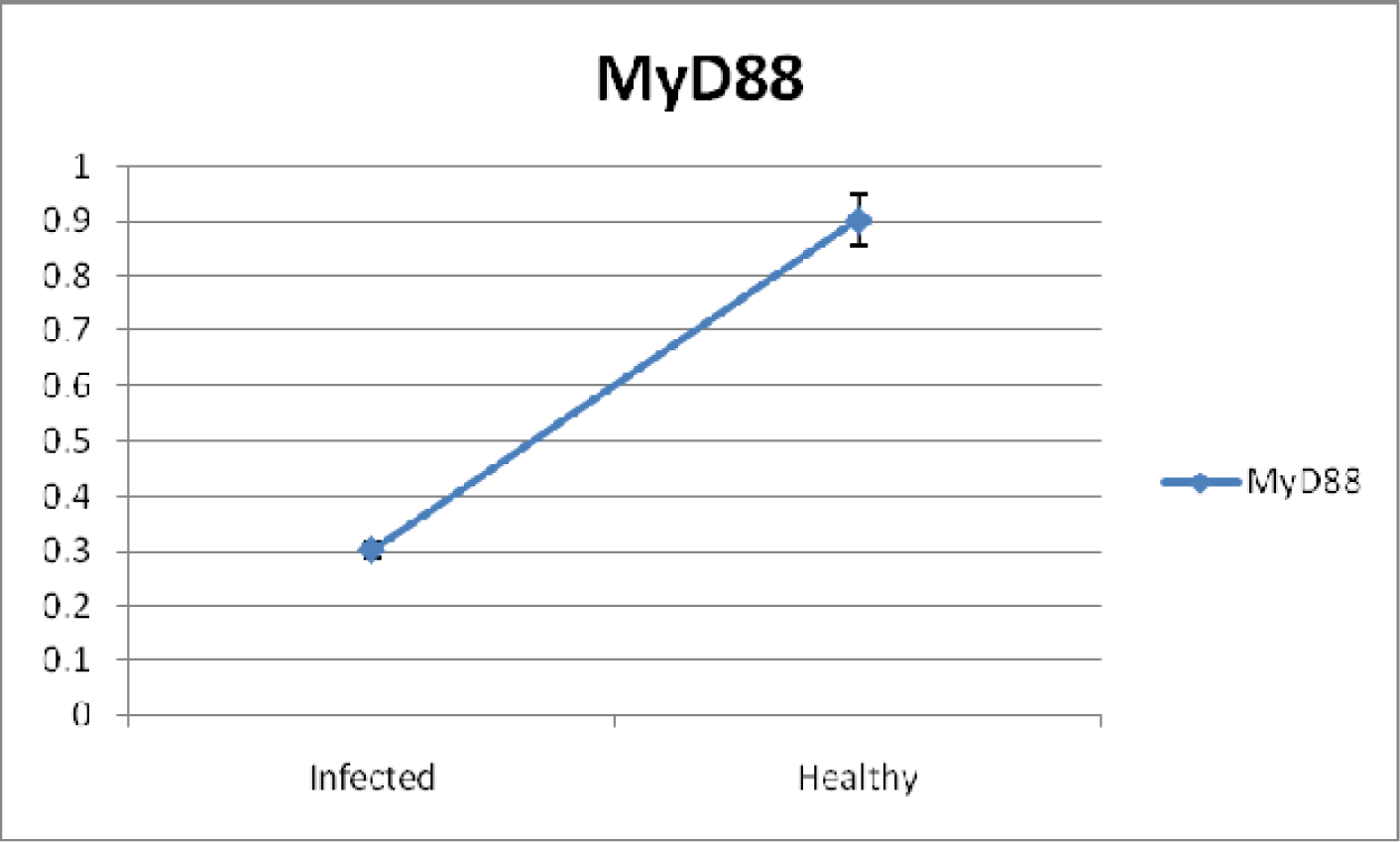
Differential mRNA expression profiling for MyD88 with respect to infected and healthy sheep with *Haemonchus contortus*.

**Fig 3:**
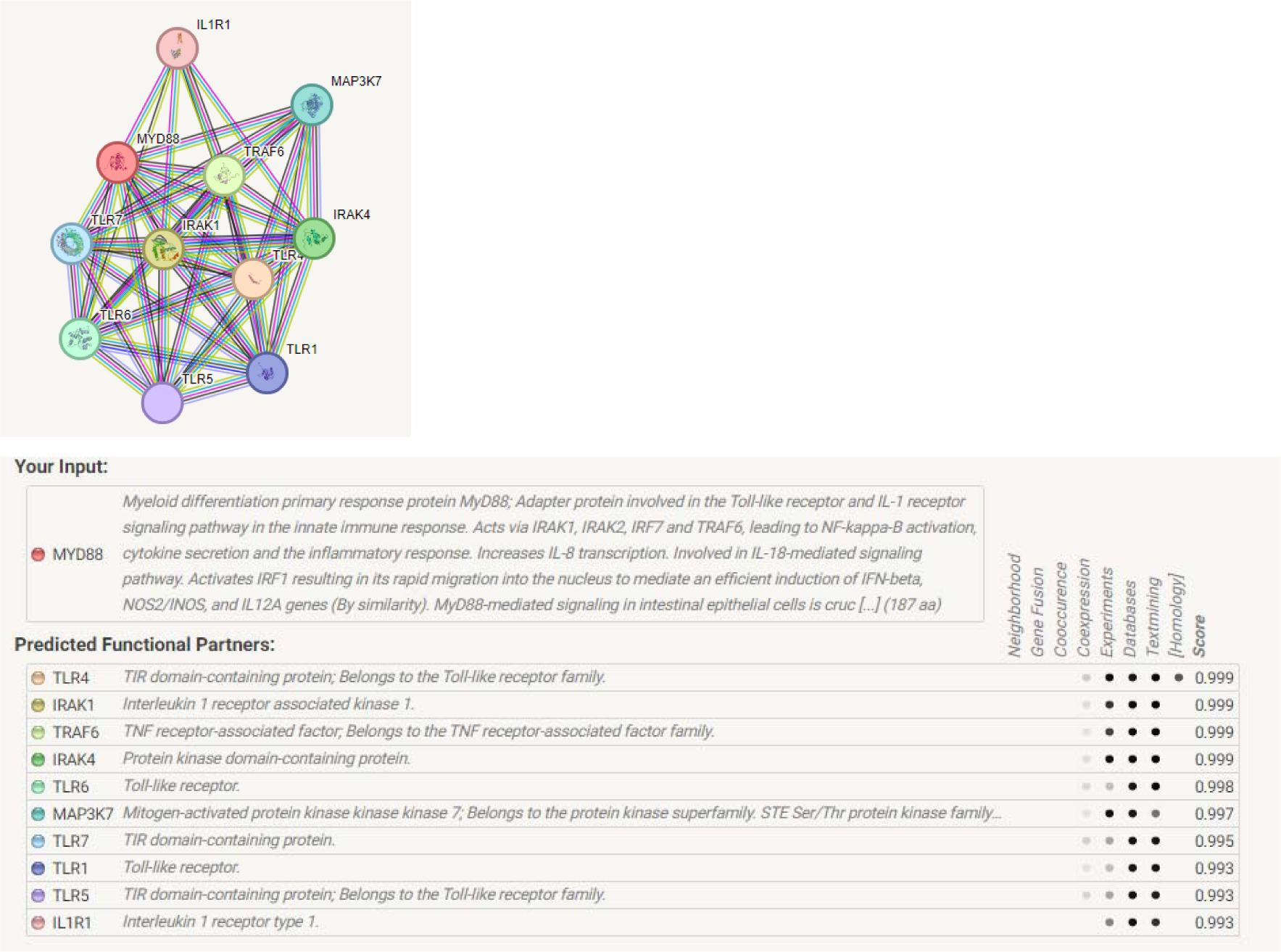
STRING Network analysis for MYD88 with other genes in sheep.

### KEGG Analysis

KEGG pathway analysis depicts the role of MyD88 through TLR signalling pathway (Fig 4a), MAPK signalling pathway(Fig 4b), NF kappa signalling pathway(Fig 4c), NOD like receptor pathway (Fig 4d). Pathway analysis depicting the role of MyD88 in resistance to other parasite, namely.

**Fig 4a:**
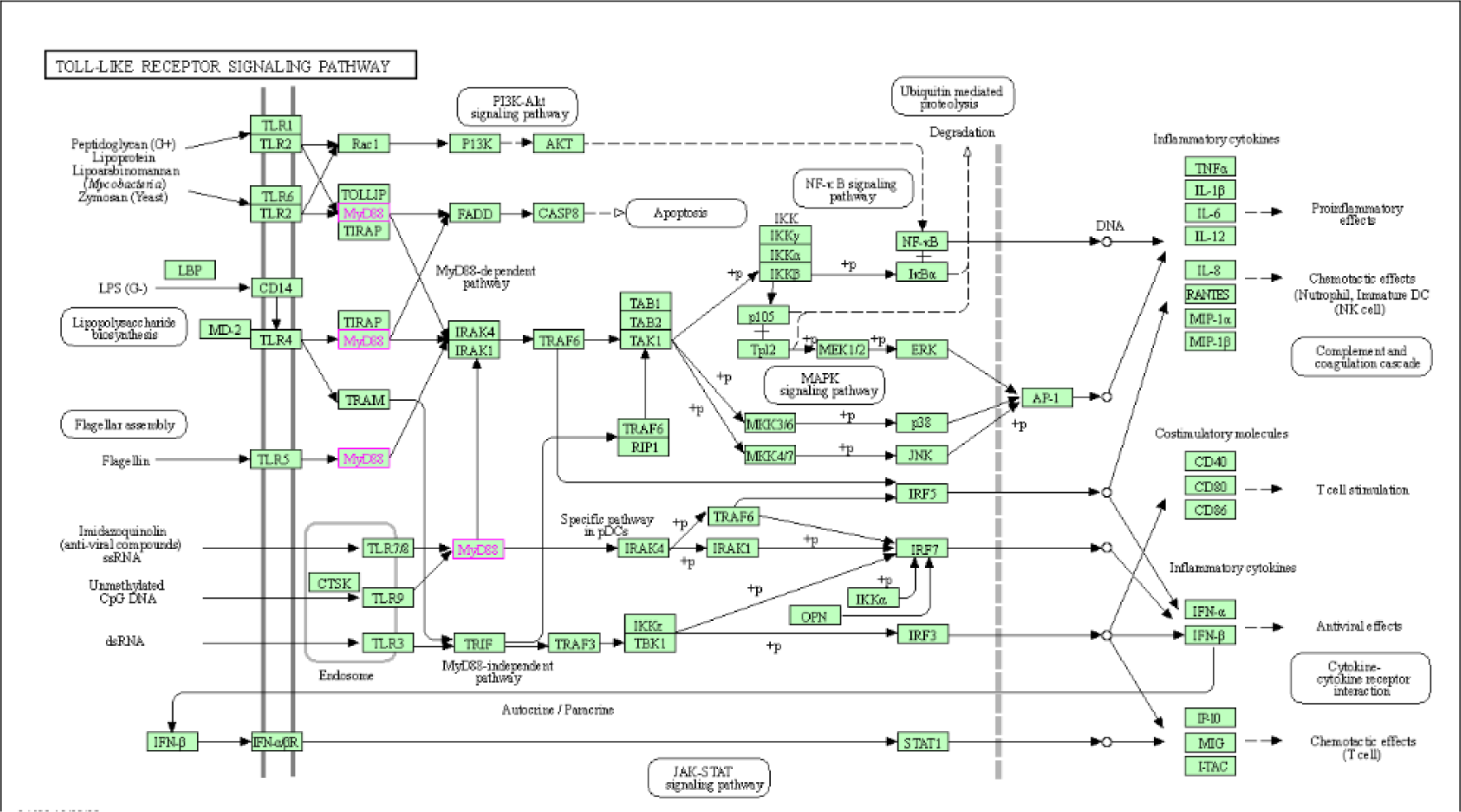
KEGG pathway analysis depicted the role of MyD88 through TLR signalling pathway.

**Fig 4b:**
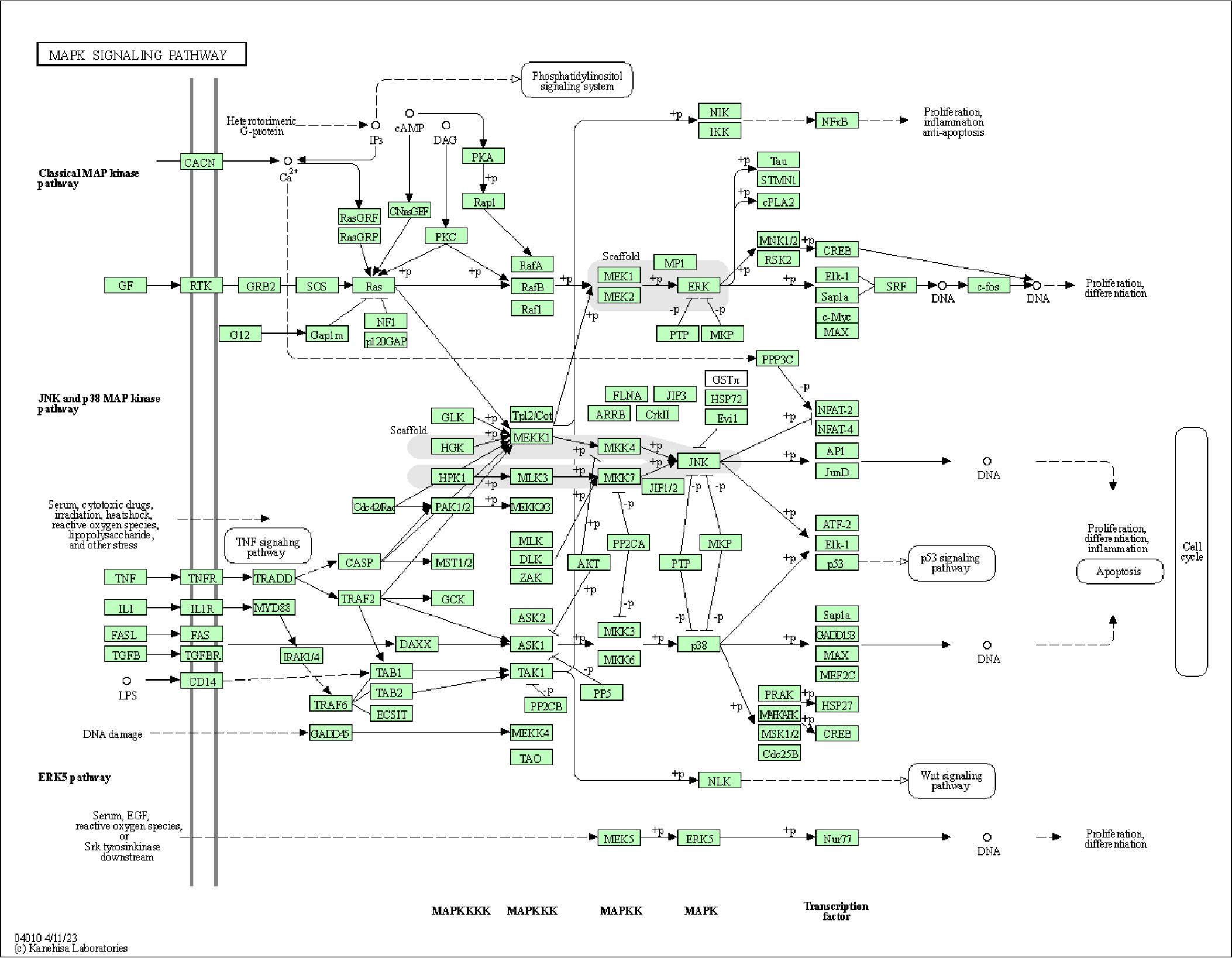
KEGG pathway analysis depicted the role of MyD88 through MAPK signalling pathway.

**Fig 4c:**
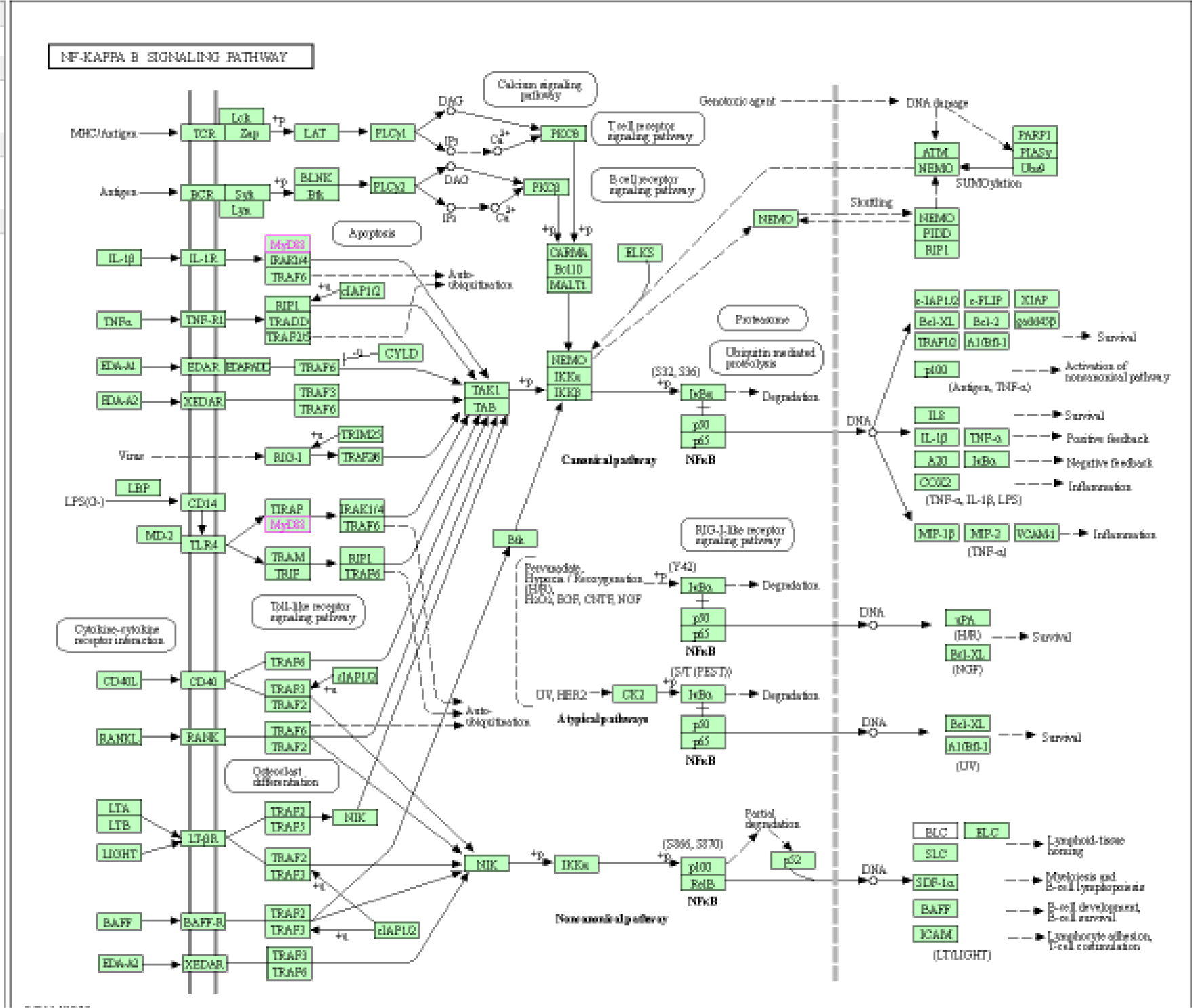
KEGG pathway analysis depicted the role of MyD88 through NF kappa signalling pathway.

**Fig 4d:**
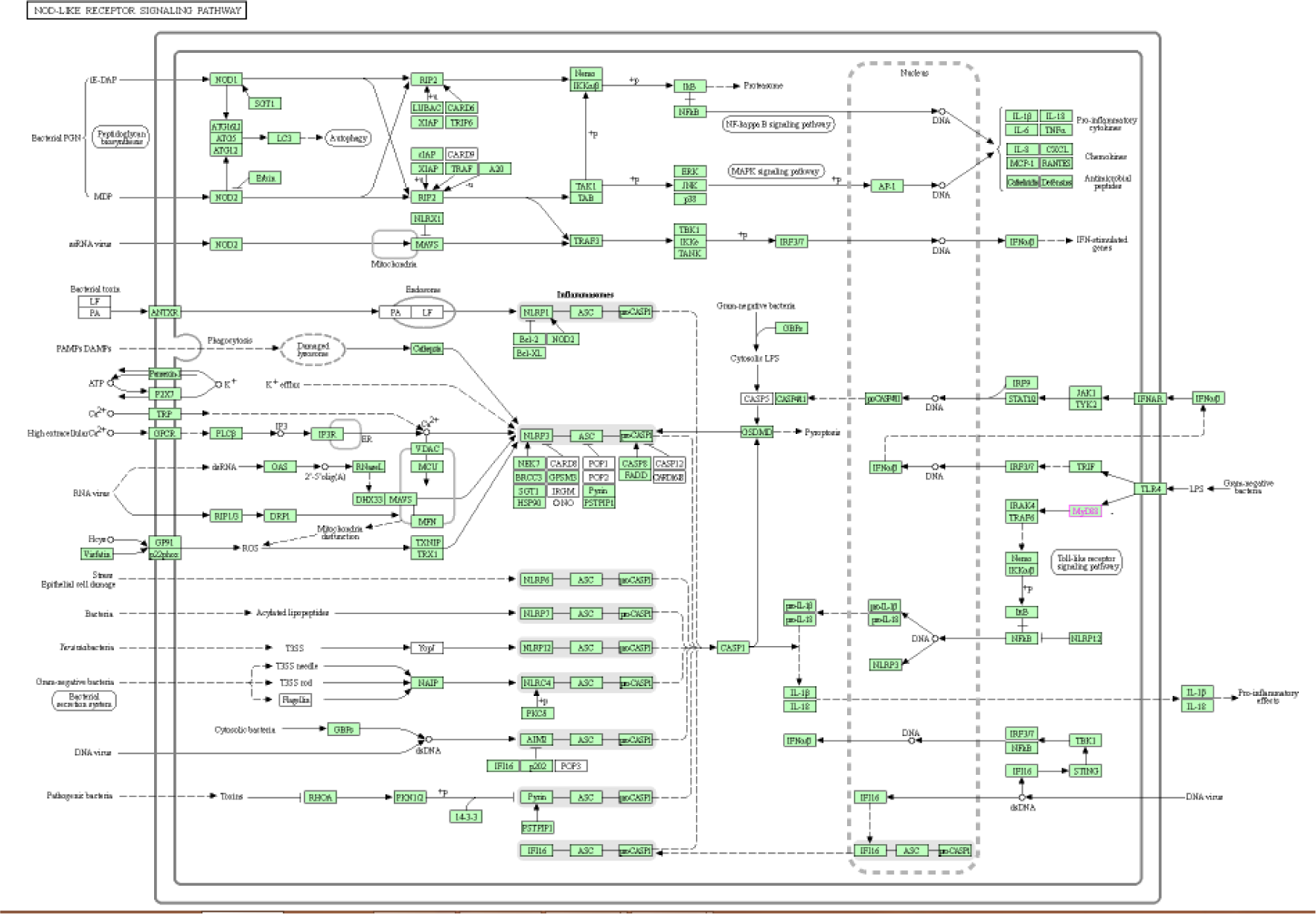
KEGG pathway analysis depicted the role of MyD88 through NOD like receptor pathway.

### Pathway analysis with other parasites

KEGG pathway analysis indicates the role of MyD88 against other diseases caused by parasites, namely, Toxoplasmosis (Fig 4e), African Trypanosomiasis(Fig 4f) and Chagus disease(Fig 4g).

**Fig 4e:**
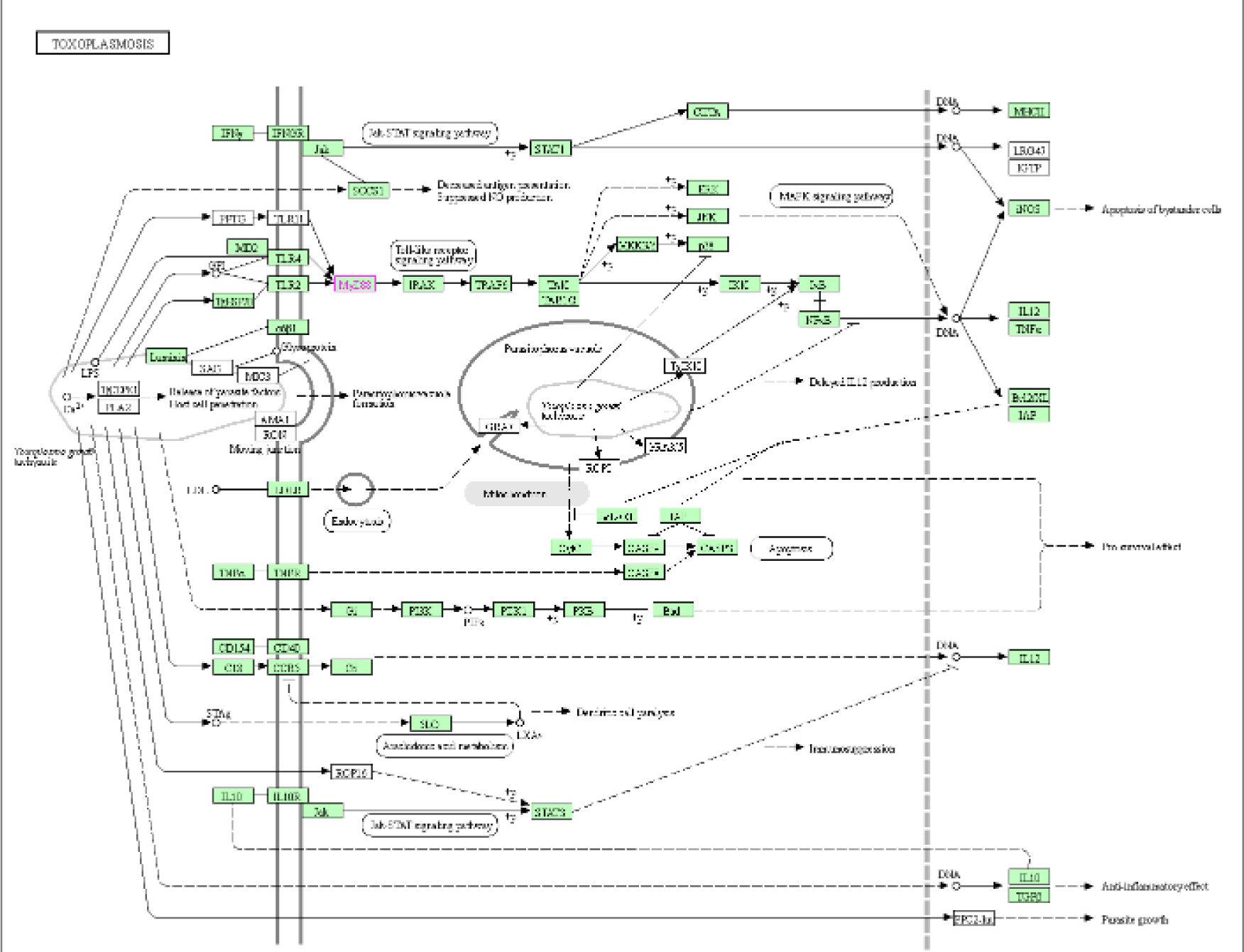
KEGG pathway analysis depicted the role of MyD88 against toxoplasmosis.

**Fig 4f:**
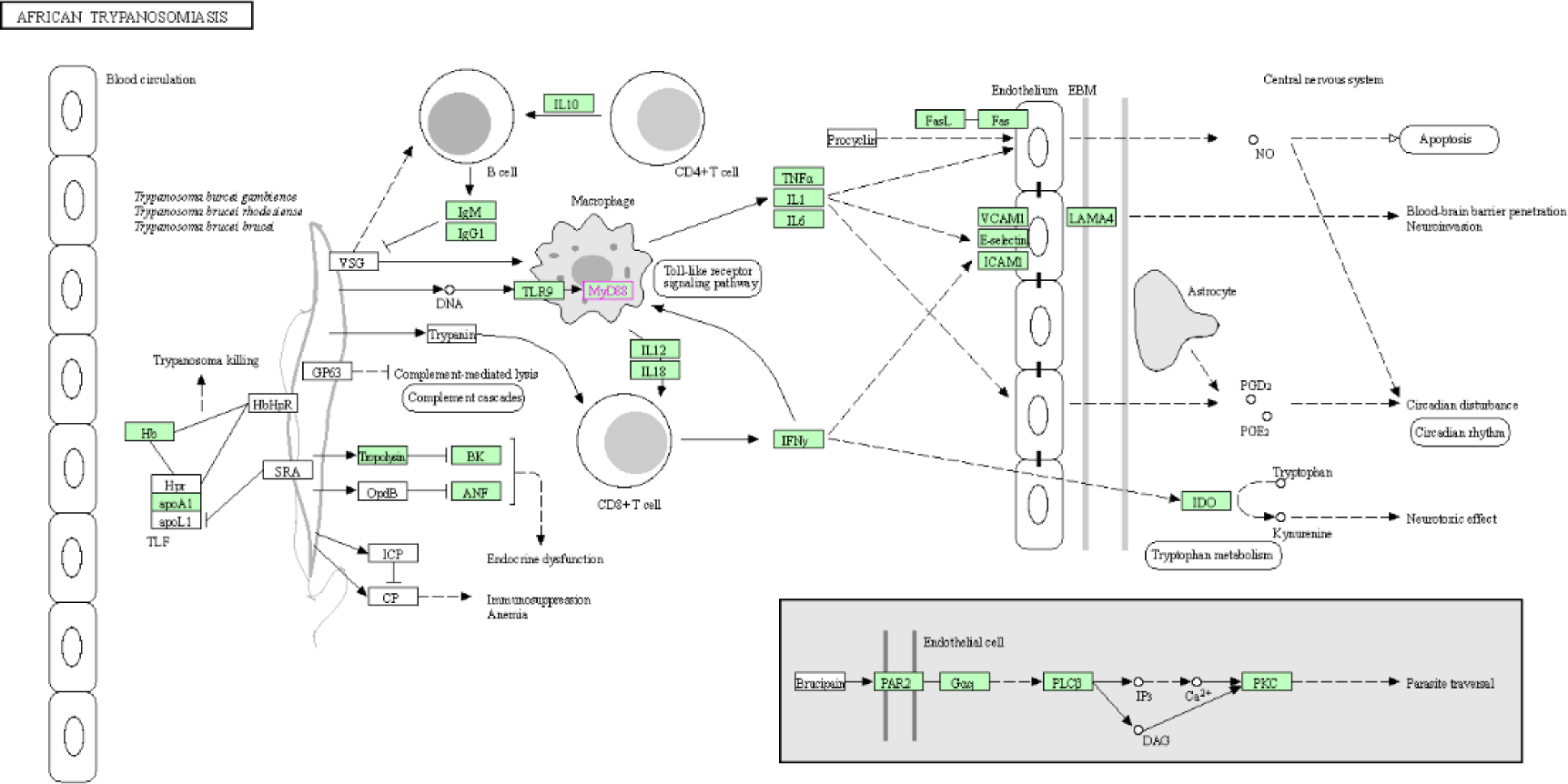
KEGG pathway analysis depicted the role of MyD88 against African Trypanosomiasis.

**Fig 4f:**
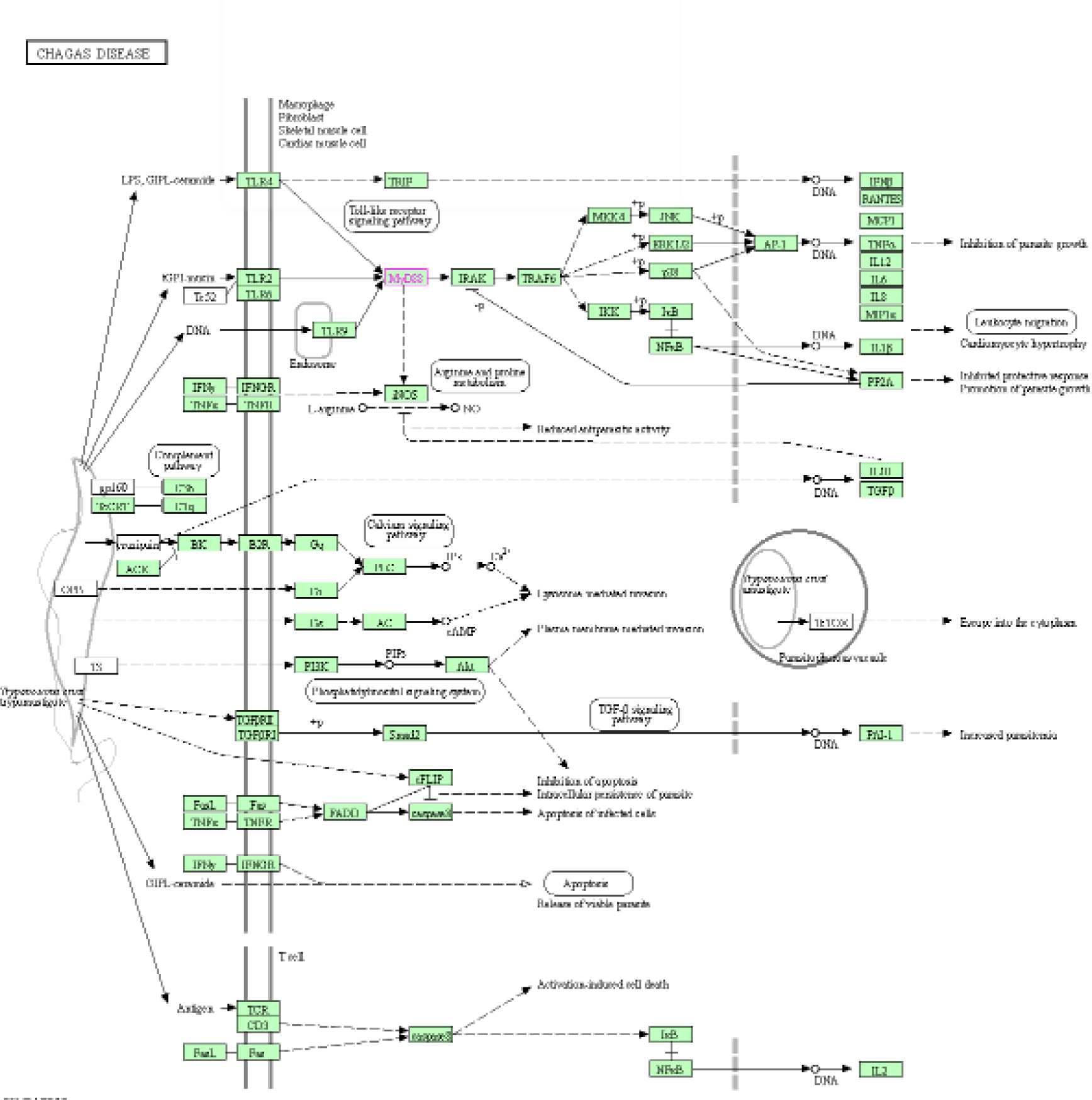
KEGG pathway analysis depicted the role of MyD88 against Chagas disease.

The current study reveals that expression of MyD88 is trigerred in sheep infected with *Haemonchus contortus* in comparison to healthy sheep.

MyD88 (Myeloid Differentiation Primary Response gene 88) transfers signals from certain proteins called Toll-like receptors and interleukin-1 (IL-1) receptors, which are important for an early immune response to foreign invaders such as bacteria. In response to signals from these receptors, the MyD88 adapter protein stimulates signaling molecules that turn on a group of interacting proteins which are nuclear factor-kappa-B. It is an important adaptor protein which is used by every Toll like receptor except TLR3. TIRAP (Toll/Interleukin-1 receptor domain containing adaptor protein) is required for the recruitment of MyD88 to TLR-2 and TLR-4 and then it plays its part and signalling occurs through IRAK (Interleukin-1 receptor associated kinase) (Arancibia SA et.al. 2007.). The MYD88 gene instructs to make a protein engaged in signalling in the immune cells. The MyD88 protein acts as an adapter, connecting proteins that receive signals from outside the cell to the proteins that relay signals inside the cell.

Myeloid Differentiation Primary response gene 88 is encoded in humans. MYD88 gene provides instructions for making a protein involved in signaling within immune cells. The MyD88 protein acts as an adapter, connecting proteins that receive signals from outside the cell to the proteins that relay signals inside the cell. In particular, MyD88 transfers signals from certain proteins called Toll-like receptors and interleukin-1 (IL-1) receptors, which are important for an early immune response to foreign invaders such as bacteria. In response to signals from these receptors, the MyD88 adapter protein stimulates signaling molecules that turn on a group of interacting proteins which are nuclear factor-kappa-B.

Some available evidences suggest that MyD88 is dispensable for human resistance against some viral infections and to all but a few pyogenic bacterial infections (Von Bernuth H, Picard C, Jin Z, Pankla R, et.al., 2008.). It has been reported that MCP1 secretion which is a chemokine required for monocyte recruitment is induced by the invasive bacteria *L. monocytogenesis* is MyD88 independent but based on this it is also evident that MyD88 deficiency does not impair monocyte recruitment to *L. monocytogenesis* infected spleens but prevent the activation of the monocyte (Natalya V. Serbina, et.al. 2003.). In a study conducted by Ives and Masina et.al., they investigated innate immune responses adapter protein modulators and showed that both MyD88 and TLR-9 played a vital role in the development of Th1 dependent healing responses against L. guyanesis parasites regardless of their LRV(Leishmania RNA Virus) status. The absence of MyD88 and or TLR-9 dependent signalling pathways resulted in increased Th2 associated cytokines (IL-4 and IL-13) which were corresponded with low transcript levels of IL-12p40. Dependency of IL-12 was further confirmed in the mice which were completely susceptible to the infection. Protection to L.guyanesis infection triggered by MyD88 and TLR-9 dependent immune responses arises independently to those induced due to high LRV burden within the parasite (Annette Ives https://journals.plos.org/plosone/article?id=10.1371/journal.pone.0096766).

The discharge of gastrointestinal nematode *T.muris* is mediated by a T helper 2 type response involving IL-4 and IL-13. In a study conducted by Helmby and Grenbis, they showed that Th1 response associated susceptibility is dependent on activation signals by MyD88 and TLR-4. Mice deficient in these two genes, are highly resistant to *T.muris* infection and develop strong antigen specific Th2 responses in mucosa associated lymphoid tissues. Here MyD88 and TLR4 not only develops pro inflammatory response against bacterial infection but also against some helminths. They developed and demonstrated first that these two genes are involved in the connection of innate and acquired immune response against chronic gastro intestinal nematodes (Helena Helmby and Richard K, 2003). Hasan and Chaffois et.al., carried out a study which revealed that TLR10 in human is a functional receptor which is expressed by B-cell and plasmacytoid dendritic cell which activates gene transcription through MyD88 (Uzma Hasan, Claire Chaffois et.al., 2005;).

Studies have shown that MyD88 or IPS 1 signalling pathway are able to control primary antiviral responses and intranasal influenza A virus infection whereas the protective adaptive immune responses influenza A virus are ruled by the TLR7-MyD88 pathway (Shohei Koyama,et.al.. 2007.). Robert O. Watson et. al., (2006) showed that mice deficient with MyD88 is stably colonized by *Campylobacter jejuni* along with this Nramp1 deficiency increases the mouse susceptibility to *C. jejuni* which is an indication that MyD88 deficient mice can be used as a model to study the colonization of *C.jejuni* an the role of Nramp1 I control of this pathogenic bacteria (Robert O. Watson, et.al. 2006.).

*Toxoplasma gondii* causing toxoplasmosis is one of the most common parasitic infection in human. It forms tissue cysts in the brain and can cause life-threatening toxoplasmic encephalitis in immune compromised patients. For the innate sensing of *Toxoplasma gondii* TLR adaptor MyD88 activation is required. Mice deficient in MyD88 gene have defective IL-12 and Th1 effector responses, and are susceptible to the acute phase of T. gondii infection. In the experiment conducted by them, MyD88-deficient mice and control mice were given oral infection of *T. gondii* cysts. Cellular and parasite infiltration in the peripheral organs and in the brain were done by histology and immune histochemistry. Cytokine levels were analysed by ELISA and chemokine mRNA levels were quantified by real-time PCR (qPCR). Thirteen days post the infection period, a higher parasite burden was observed but there was no histological change in the liver, heart, lungs and small intestine of MyD88 -/- and MyD88 +/+ mice. But MyD88 mice compared to MyD88 +/+ mice were highly susceptible to cerebral infection, showed high parasite load in the brain, severe neuropathological signs of encephalitis and capitulated within 2 weeks of oral infection. Susceptibility was primarily associated with lower expression of Th1 cytokines, especially IL-12, IFN-γ and TNF-a,decrease in the expression of CCL3, CCL5, CCL7 and CCL19 chemokines, marked defect of CD8 + T cells, and infiltration of CD11b + and F4/80 + cells in the brain. So it was concluded by them that MyD88 is essential for the protection of mice during the cerebral installation of *T. gondii* infection. These results establish a role for MyD88 in T cell-mediated control of *T. gondii* in the central nervous system (CNS) (Marbel Torres, et.al.2013,)

Double stranded RNA Bluetongue virus induces Type 1 interferon in Plasmacytoid dendritic cells by MyD88 dependent and TLR 7/8 independent signaling pathway (Suzana Ruscanu, Florentina Pascale et.al., 2012.). MyD88 knockout mice have shown significant increase in *S.aureus* biofilm and further immunofluorescence staining of biofilm infected tissues result in fibrosis (Hanke ML et.al. 2012). Libo Su et.al., carried out the experiment to determine how the helminth parasite, *H.polygurus* affect the TLR signalling pathway during the time of its reponse in an enterobacteria. MyD88 mice and a knockout mice that is the wild type C57BL/6 mice were infected with *H.polygyrus* and *Citobacter rhodentium* or both. They observed that the knockout mice coinfected with both parasitic and helmintic infection developed severe intestinal inflammation and higher chances of mortality than the wild type mice. The increased susceptibility to *C.rodentium*, intestinal injury and mortality of the coinfected MyD88 knock out mice were observed to be correlated with reduced intestinal phagocyte recruitment and an increase in bacterial translocation. The increase in the bacterial disease and infection severity were found to be associated with a downregulation of anti microbial peptide expression in the intestinal tissue in co-infected MyD88 knockout mice.

Their results suggested that the MyD88 signalling pathway plays a critical role for host defense and survival during helminth and enteric bacterial co-infection(LiboSu,et.al.Shihttps://journals.plos.org/plosntds/article?id=10.1371/j ournal.pntd.0002987) MyD88 gene, as we can see in the figure, contrastingly shows the over expression in the healthy sheep and it is under expressed in diseased sheep. Several studies have been made to demonstrate that MyD88 associated with TLR9 played an important role in the development of the Th1 dependent responses for healing against L.guyanensis parasite nevertheless of the Leishmania RNA virus status. Th2 associated cytokines (Il-4, IL-13) are increased in the absence of MyD88-TLR9 dependent signalling pathway (Ives A, et al. (2014)). Studies by Helmby and Grencis states that MyD88 helps in traversing the innate and acquired immune response in a gastrointestinal nematode. TLR4 and MyD88 deficient mice are resistant to chronic T.muris and acquire strong antigen specific Th2 response in mucosa associated lymphoid tissue. Therefore these genes play a pivotal role not only in the pro inflammatory responses but also in the responses against helminths and parasites. They experimentally stated that MyD88 is required for the development of atrocious gastrointestinal nematode infection. MyD88 Knockout mice secrete high levels of Th2 cytokines in the course of infection by T.muris (Helena Helmby and Richard K. Grencis. 2003).

Su et.al., highlighted an important global issue which is the infection of helminth along with the co infection of bacterial pathogen like Escherichia coli. Scientists stated that MyD88 kockout mice along with the co-infection of H.polygyrus and C.rodentium matured into more acute intestinal inflammation and increased mortality compared to the wild type mice. The susceptibility to C.rodentium, intestinal injury and and mortality of co-infected MyD88 Knockout mice were found to be associated with the noticeable reduction in the recruitment of intestinal phagocyte. Along with this, the increase in bacterial infection, severed disease infection is co-related with a down regulation of anti microbial peptide expression in the intestinal tissue of the co-infected MyD88 KO mice and therefore they suggested that MyD88 signalling pathway plays a pivotal role for host defense and survival during helminth and bacterial co-infection (Su L, Qi Y, et al. (2014)).

Reynolds et.al., also studied that the MyD88 signalling hinders the protective immunity to the Gastrointestinal Helminth Parasite. Heligmosomoides polygyrus. In the study, they investigated the result of a primary H. polygyrus infection in MyD88 signaling deficient mice and found that they were more resistant to infection than wild-type C57BL/6 mice and had a higher frequency of IL-4–producing CD4+ T cells post infection. Furthermore, MyD88-deficient mice evolved with large numbers of granulomas throughout the intestinal wall in response to the infection. Hence they concluded that the absence of MyD88 signaling resulted in elevated H. polygyrus ejection, a phenotype that was partially replicated in TLR2-deficient mice, but not in mice deficient in TLR4, TLR5, or TLR9 which gave rise to two possibilities: that signals through individual TLRs redundantly maintain helminth susceptibility or that a TLR independent pathway is key to the elevated immunity of MyD88 deficient mice (Lisa A. Reynolds et.al., 2014;)

### Conclusion

MyD88 has proved to have an important role in providing immunity against *Haemonchus contortus*

## Acknowledgement

The authors are thankful to Department of Biotechnology, Ministry of Science and Technology, Govt. of India (Grant number BT/PR24310/NER/95/649/2017) and Department of Science and Technology, Govt. of India (Grant no. EMR/2016/003554) for providing the financial support. The technical and financial support by Vice-Chancellor, West Bengal University of Animal and Fishery Sciences is duly acknowledged. Thanks to Director, AH & VS, Animal Resource Development Department, Govt. of West Bengal.

---

**SupplementaryFig 1:**
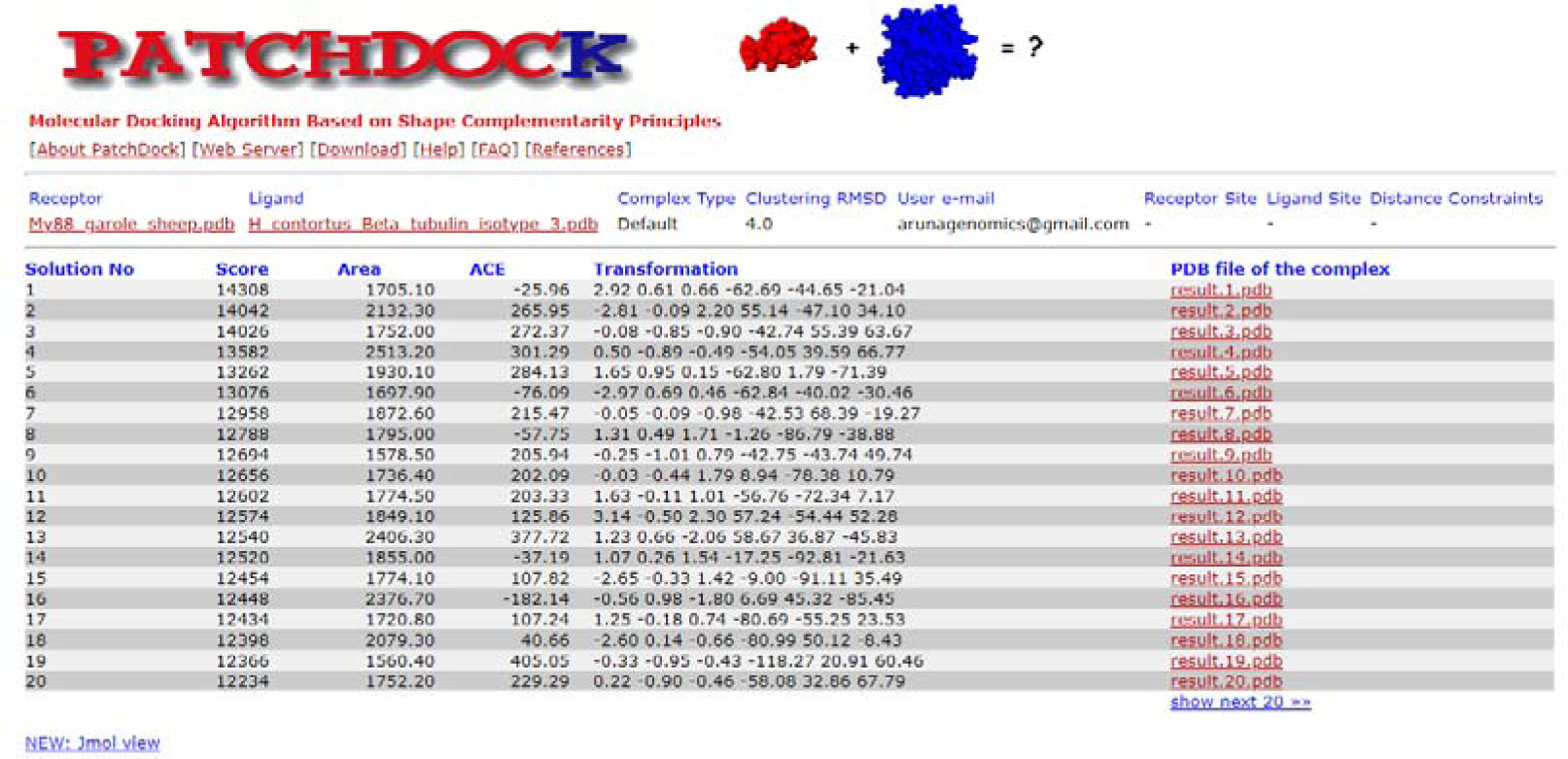
Patchdock score for MyD88 with Beta tubulin surface protein of *Haemonchus contortus*.

**SupplementaryFig 2:**
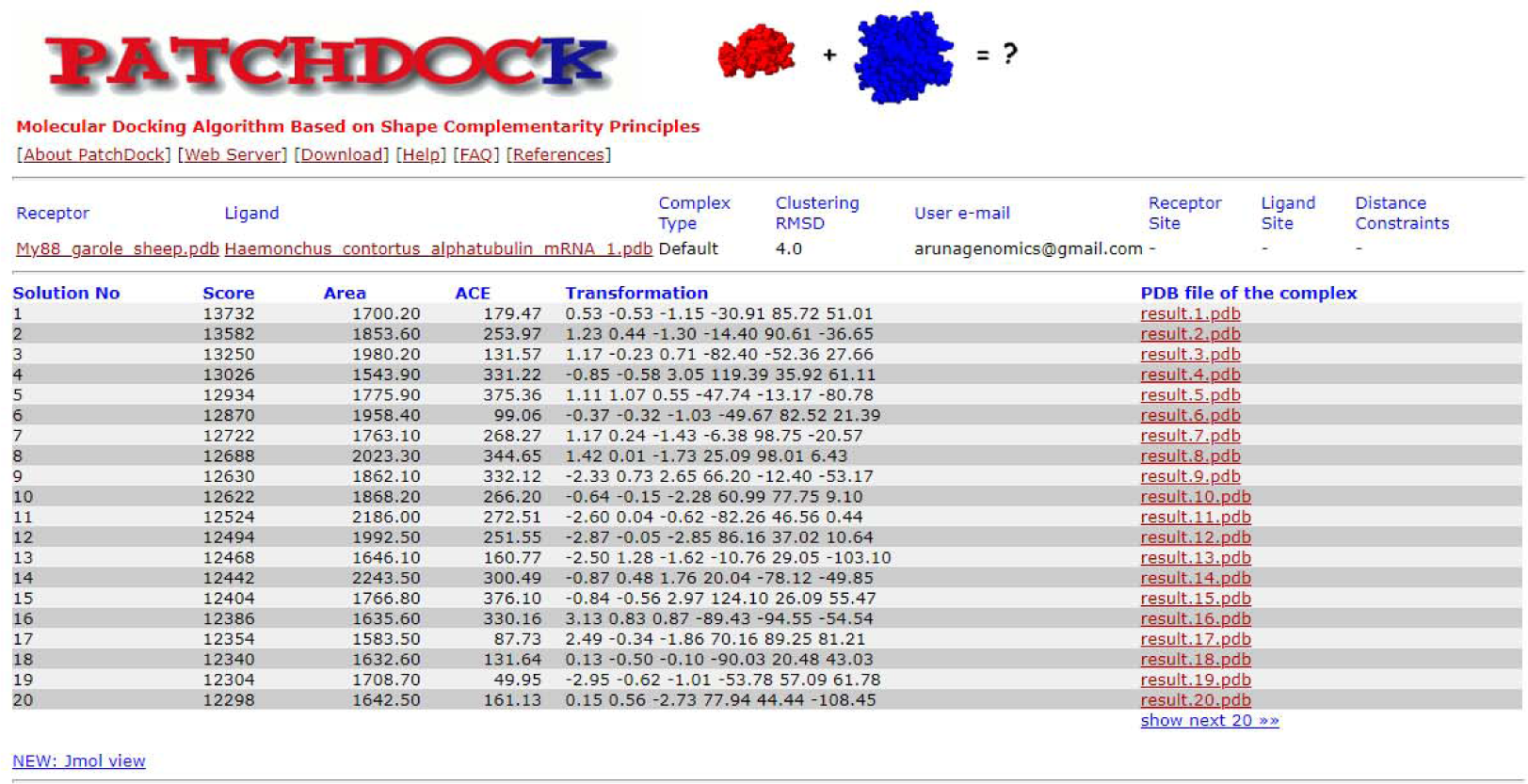
Patchdock score for MyD88 with alpha tubulin surface protein of *Haemonchus contortus*.

